# Induced pluripotent stem cell-derived astrocytes from patients with schizophrenia exhibit an inflammatory phenotype that affects vascularization

**DOI:** 10.1101/2022.03.07.483024

**Authors:** Pablo Trindade, Juliana M. Nascimento, Bárbara S. Casas, Tomás Monteverde Faúndez, Juciano Gasparotto, Camila Tiefensee Ribeiro, Sylvie Devalle, Daniela Sauma, José Claudio Fonseca Moreira, Daniel Pens Gelain, Lisiane O. Porciuncula, Verónica Palma, Daniel Martins-de-Souza, Stevens Kastrup Rehen

## Abstract

Molecular and functional abnormalities of astrocytes have been implicated in the etiology and pathogenesis of schizophrenia (SCZ). In this study, we examined the proteome, inflammatory responses, and secretome effects on vascularization of human induced pluripotent stem cell (hiPSC)-derived astrocytes from patients with SCZ. Proteomic analysis revealed alterations in proteins related to immune function and vascularization. Reduced expression of the nuclear factor kappa B (NF-κB) p65 subunit was observed in these astrocytes, with no incremental secretion of cytokines after tumor necrosis factor alpha (TNF-α) stimulation. Among inflammatory cytokines, secretion of interleukin (IL)-8 was found particularly elevated in SCZ-patient-derived– astrocyte conditioned medium (A_SCZ_CM). In a chicken chorioallantoic membrane (CAM) assay, A_SCZ_CM reduced the diameter of newly grown vessels, and this effect could be mimicked with exogenous addition of IL-8. Taken together, our results suggest that SCZ astrocytes are immunologically dysfunctional and may consequently affect vascularization through secreted factors.

## Introduction

Schizophrenia (SCZ) is a psychotic disorder characterized by the manifestation of a wide range of behavioral and cognitive symptoms that affects around 0.5–1% of the population worldwide. Despite its relatively high frequency, the early pathophysiology of this mental illness is still not well characterized ^1^. Diagnostically, the term SCZ is an umbrella term that encompasses a spectrum of presentations with overlapping behavioral or psychiatric changes inclusive of different degrees of severity ^2^. The heterogeneity of clinical symptoms associated with SCZ suggests that the etiology and course of the disorder may be influenced by environmental, psychosocial, and genetic factors ^3^.

Several biological mechanisms have been related to the onset of SCZ symptoms, including neurodevelopmental impairments ^4^, disruptions in neurotransmitter systems ^5^, and multigenic risk factors ^6^. Immune dysfunctions in patients with SCZ have also offered insights into potential etiological mechanisms. Indeed, there is a growing body of recent evidence supporting the notion that SCZ may be a disorder of immune responses within the brain ^7, 8^, with glial cells likely playing a major role.

Astrocytes and microglia are the major immune response modulators of the central nervous system (CNS) and they exert effects via secreted cytokines ^9, 10^. Beyond their well-documented roles in synaptic modulation and energy metabolism, astrocytes near capillaries act as an important component of the blood brain barrier (BBB) by modulating brain vascularization and serving as an important source of inflammatory cytokines ^11^. Astroglial alterations observed in SCZ have mostly been focused on astrocyte-mediated disruption of neurotransmission ^12, 13^. Meanwhile, the astroglial immune response in SCZ remains poorly characterized.

Inflammation has been observed in both neurodegenerative and psychiatric disorders, and the astroglial changes that have been described in SCZ specifically are not uniform and often divergent. For instance, while elevated levels of reactive astrogliosis have been reported for patients with SCZ in some reports ^14–16^, others observed no change or even a decrease in the same parameters ^17–19^. This heterogeneity could be explained by differences in the brain regions analyzed, SCZ stage, age of patients, ongoing treatment, and severity of symptoms. Both vascular abnormalities and increased circulating cytokines have been reported systematically in SCZ, but the role astrocytes play in modulating these events is unknown ^20, 21^.

Postmortem proteomic analysis of selected brain area samples from patients with SCZ, including prefrontal cortex ^22^, corpus callosum ^23, 24^, and hippocampus ^25^ samples, have revealed alterations in two common astrocyte markers, namely aldolase C (ALDOC) and glial fibrillary acidic protein (GFAP) ^26, 27^. Higher levels of S100 calcium-binding protein β (S100β) were found in the brains of patients who had paranoid schizophrenia than in the brains of patients who had residual SCZ ^28^, suggesting that inflammation may be associated with distinct symptoms ^29, 30^. However, despite the extensive literature addressing neurophysiological aspects of SCZ spectrum disorders, there have been few studies that have focused on cell-specific and astroglial alterations in SCZ.

Advances in the development of patient-derived induced pluripotent stem cells (iPSCs) have shed light on the impact of immune dysfunction in relation to SCZ onset ^31^. Studies using human iPSCs (hiPSCs) or hiPSC-derived neural cells from patients with SCZ showed that high levels of reactive oxygen species were associated with the presence of SCZ-related behavioral ^32–35^. Additionally, hiPSC-derived glial progenitors from patients with SCZ showed delayed astroglial differentiation and abnormal migration into the cortex after being administered as chimeric implantations in mice ^36^. When hiPSC-derived astrocytes from SCZ patients were exposed to interleukin (IL)-1β, disturbances in signaling pathways related to T cell recruitment were observed ^37^.

In a previous study, our group showed that angiogenesis was affected by conditioned media from neural stem cells (NSCs) derived from the hiPSCs of patients with SCZ ^38^. Additionally, we showed recently that hiPSC-derived astrocytes stimulated with tumor necrosis factor alpha (TNF-α) can reproduce canonical events associated with reactive astrogliosis ^39^. Building from these prior studies, the aim of the present study was to examine the proteome, inflammatory responses, and secretome effects on vascularization of hiPSC-derived astrocytes from patients with SCZ. We compared SCZ-patient-derived–astrocyte conditioned medium (A_SCZ_CM) and control-subject-derived–astrocyte conditioned medium (A_CON_CM) in chicken chorioallantoic membrane (CAM) assays to evaluate the impact of cytokines secreted from cultured hiPSC-derived astrocytes from SCZ patients on angiogenesis. We further compared the CAM results obtained with A_SCZ_CM to those obtained with exogenous cytokine addition to test the hypothesis that a cytokine may have a vascularization disrupting effect in the context of SCZ.

## Materials and Methods

### Cell lines and generation of hiPSC-derived astrocytes

Human astrocytes were differentiated from hiPSC-derived NSCs obtained from the iPSCs of four healthy subjects and three patients diagnosed with schizophrenia-spectrum disorder. These cell lines have been characterized previously ^38, 39^. Four control subject cell lines were used: one from the Coriell Institute Biobank (Subject 1) and the other three from subjects whose cells were reprogrammed at the D’Or Institute for Research and Education (Subjects 2, 3, and 4). To decrease genetic heterogeneity among SCZ cell lines, two of the SCZ patients were siblings (Patients 1 & 2); their cells were purchased from the Coriell Institute Biobank. The third SCZ cell line was reprogrammed at the D’Or Institute for Research and Education (Patient 3). Reprogramming of human cells was approved by the ethics committee of Copa D’Or Hospital (CAAE number 60944916.5.0000.5249, approval number 1.791.182). Human cell experiments were performed according to Copa D’Or Hospital regulations. The identities and properties of the cell lines used are summarized in Supplementary Table 1.

The hiPSC-derived astrocytes used in this study were cultured as described previously ^39^. Briefly, NSCs were seeded at a density of 5 × 10^4^ cells/cm^2^ and precoated with Geltrex (A1413301, Thermo Fisher Scientific) in NSC expansion medium. The next day, the medium was replaced with astrocyte induction medium constituted by DMEM/F12 (11330-032, Thermo Fisher Scientific), 1× N2 supplement (17502001, Thermo Fisher Scientific), and 1% fetal bovine serum (12657029, Thermo Fisher Scientific). Astrocyte induction medium changes were performed every other day for 21d. From then on, cells were kept in culture for an additional 4 weeks to allow expansion and maturation of astroglial functions. All experiments described in this manuscript were performed at least 49 d after the beginning of astrocyte differentiation.

### Nuclear factor kappa B (NF-κB) translocation experiments and immunocytochemistry

For astroglial markers labeling, cultured hiPSC-derived astrocytes were seeded at a density of 6.25 × 10^3^ cells/cm^2^ in 24-well plates with treated glass coverslips. For the NF-κB translocation experiments, 96-multiwell cell-carrier plates were used (PerkinElmer, USA). Four days after plating, multiwell-plated cells were serum-deprived and, on the following day, TNF-α (1–50 ng/ml) was added for 1 h. The cultures were fixed with 4% paraformaldehyde in phosphate-buffered saline for 20 min, permeabilized with 0.3% Triton X-100, and exposed to blocking solution containing 2% bovine serum albumin or 5% normal goat serum (all from Sigma Aldrich, USA). The fixed cells were incubated overnight at 4 °C with the following primary antibodies: mouse anti-GFAP (1:200; MO15052, Neuromics, Minneapolis, MN), rabbit anti-ALDH1L1 (1:1000; ab190298, Abcam, Cambridge, UK), mouse anti-vimentin (1:500; MO22179, Neuromics, Minneapolis, MN), rabbit anti-EAAT1 (1:100; ab416, Abcam, Cambridge, UK), rabbit anti-EAAT2 (1:200; ab41621, Abcam, Cambridge, UK), rabbit anti-connexin 43 (1:100; #3512, Cell Signaling, USA) and mouse anti-NF-κB p65 (1:100; sc-8008, Santa Cruz Biotechnology, USA). Subsequently, the samples were incubated with the following secondary antibodies (all from Thermo Fisher Scientific, USA): goat anti-mouse Alexa Fluor 488 IgG (1:400; A-11001), goat anti-rabbit Alexa Fluor 594 IgG (1:400; A-11037), and goat anti-mouse Alexa Fluor 594 IgG (1:400; A-11032). Cell nuclei were stained with 0.5 μg/ml 4′-6-diamino-2-phenylindole for 5 min. Astroglial marker labeling images were acquired on a TCS-SP8 confocal microscope (Leica, Wetzlar, Germany). NF-κB translocation images were acquired via in an Operetta high-content imaging system (PerkinElmer, USA) with a 20× objective. The total number of cells was calculated based stained nuclei counts. Densitometry analyses were performed in Harmony 5.1 software (PerkinElmer, USA). Each experimental condition was represented by eleven distinct fields from wells in triplicate.

### Real-time quantitative polymerase chain reaction (RT-qPCR)

Cultures of hiPSC-derived astrocytes were seeded at a density of 2 × 10^4^ cells/cm^2^ in T-75 culture flasks. Four days after plating, cells were harvested with Accutase (Merck, Darmstadt, Germany) and centrifuged at 300 ×*g* for 5 min. The supernatant was discarded and the pellet was submitted to RNA extraction. Total RNA was isolated with TRIzol™ reagent according to the manufacturer’s instructions (Thermo Fisher Scientific, USA). RNA samples were resuspended in a final volume of 12 μl of nuclease-free water and quantified by absorbance at 260 nm in a spectrophotometer (NanoDrop 2000, Thermo Fisher Scientific, USA). One microgram of total DNase-treated RNA was reverse-transcribed in a 20-μl reaction volume using M-MLV (Thermo Fisher Scientific, USA). The resulting cDNA was diluted five-fold and RT-qPCR was performed with Eva™ Green PCR Master Mix (Biotium, Fremont, CA) in a StepOne Plus Real-Time PCR Platform (Applied Biosystems, Waltham, MA) to detect transcripts of the following genes: *NFKB1* (forward: CCC ACG AGC TTG TAG GAA AGG; reverse: GGA TTC CCA GGT TCT GGA AAC); *GAPDH*, which encodes glyceraldehyde-3-phosphatedehydrogenase (forward: 5’-GCC CTC AAC GAC CAC TTT G-3’; reverse: 5’-CCA CCA CCC TGT TGC TGT AG-3’); and *HPRT1*, which encodes hypoxanthine phosphoribosyl transferase 1 (forward 5’-CGT CGT GAT TAG TGA TGA ACC-3’; reverse: 5’-AGA GGG CTA CAA TGT GAT GGC-3’). The primers used in this study were validated by serial dilution allowing reaction efficiency calculation. Efficiencies from all tested primers ranged from 90% to 110%, not separated by >10%. RT-qPCR data were calculated by the 2^−ΔΔCt^ method using the mean values obtained for *HPRT1* and *GAPDH* to normalize the results.

### Multiplex analysis of cytokines

Cultures of hiPSC-derived astrocytes were seeded at a density of 2 × 10^4^ cells/cm^2^ in 75-cm^2^ culture flasks. Four days after seeding, cells were serum-deprived for 24 h. Next, 10 ng/ml TNF-α or vehicle was added for an additional 24 h. A_SCZ_CM samples were collected, aliquoted, and stored at −80 °C. ProcartaPlex™ bead-based multiplex immunoassays were carried out for simultaneous detection and quantitation of multiple protein targets in the A_SCZ_CM. The platform MAGPIX™ was used for simultaneous detection of IL-1β, IL-2, IL-4, IL-6, IL-8, TNF-α, IL-10, IL-13, interferon (INF)-γ, and brain-derived neurotrophic factor (BDNF) in a single sample according to the manufacturer’s instructions. Briefly, 50 μl of coated magnetic beads solution (Luminex™ Corporation) was added to each well of a 96-well plate. The loaded plates were placed onto a magnetic plate washer; the beads were allowed to accumulate on the well bottoms and then washed. We added 50-μl sample aliquots or standards to the wells and left them to incubate for 2 h at room temperature (RT) on a Capp 18-X shaking platform (500 rpm). The plates were washed and detection antibody mixture was added and incubated with the beads at RT for 30 min on the shaking platform (500 rpm). Streptavidin solution was added to each well and incubated for 30 min (500 rpm) at RT. After washing, reading buffer was added into the wells and incubated for 5 min (500 rpm) at RT. Data were acquired with MAGPIX™.

### Sample preparation for proteomic analyses

Proteomic analyses were performed with: astrocytes derived from hiPSCs from patients with SCZ; astrocytes derived from control subjects; media conditioned with TNF-α stimulated cells; and media conditioned with non-stimulated cells exposed to only vehicle. Cells were collected from hiPSC-derived astrocyte cultures seeded at a density of 2 × 10^4^ cells/cm^2^ in 75-cm^2^ culture flasks. Four days after plating, cells were harvested with Accutase (Merck, Darmstadt, Germany) and centrifuged at 300 ×*g* for 5 min. The supernatant was discarded. The pellet was washed with phosphate buffered saline and stored at −80 °C until used. Cell pellets were homogenized in lysis buffer (7 M urea, 2 M thiourea, 1% CHAPS, 70 mM DTT, and Roche EDTA-free complete protease inhibitor cocktail) for 2 h in ice, lysates were centrifuged at 10,000 ×*g* at 4 °C, and supernatant samples were collected. Initial protein content was quantified using the BCA method according to the manufacturer’s protocol (Bio-Rad Laboratories, USA). Samples (50 μg) were then subjected to sodium dodecyl sulphate– polyacrylamide gel electrophoresis, and gel slicing followed by *in-gel* reduction, alkylation, and overnight trypsin digestion (1:100, w/w) at 37 °C. The peptides obtained were extracted with 50 mM AmBic buffer and dried with a SpeedVac (Thermo Fisher Scientific, USA) and stored at −80 °C prior to quantitative and qualitative analyses by shotgun mass spectrometry (MS). Biological triplicates of both control and SCZ hiPSC-derived astrocytes were analyzed.

For conditioned media sample preparation, cultures of hiPSC-derived astrocytes were seeded at a density of 2 × 10^4^ cells/cm^2^ in 75-cm^2^ culture flasks. Four days after seeding, the cells were serum-deprived for 24 h. Next, 10 ng/ml TNF-α or vehicle was added for an additional 24 h. The conditioned media were collected, aliquoted, and stored at −80 °C until use. Conditioned media were concentrated 10-fold with Amicon Ultra centrifugal filters 3,000 MWCO (Merck-Milipore, Germany). The recovered protein was quantified with the BCA method, and 100-μg samples were subjected to gel electrophoresis and protein digestion as described above. Five conditioned media biological replicates of were analyzed for each group.

### Liquid chromatography (LC)-MS

Proteomic analyses were performed using a reverse-phase LC Acquity UPLC M-Class System coupled to a Synapt G2-Si mass spectrometer (both from Waters Corporation, MA, USA). Data were acquired by data-independent acquisition with ion mobility separation and high-definition data-independent MS. hiPSC-derived astrocyte whole-cell contents were subjected to two-dimensional LC-MS/MS analysis; for conditioned medium, first-dimension chromatography was skipped and the samples were directed to the second-dimension column. Peptides loads were separated in first-dimension chromatography on an M-Class BEH C18 column (130 Å, 5 μm, 300 μm × 50 mm, Waters Corporation, Milford, MA). Three discontinuous fractionation steps were performed (13%, 18%, and 50% acetonitrile). Peptide loads were directed, after each step, to second-dimension chromatography on a nanoACQUITY UPLC HSS T3 column (1.8 μm, 75 μm × 150 mm; Waters Corporation, USA). For whole-cell analyses, peptides were then eluted directly into a Synapt G2-Si MS platform with a 7–40% (v/v) acetonitrile gradient for 36 min at a flow rate of 0.4 μl/min. Conditioned medium samples (directed to the second-dimension column as specified above) were eluted for 95 min at a flow rate of 0.4 μl/min directly into a Synapt G2-Si MS platform. MS/MS analyses were performed by nano-electrospray ionization in positive ion mode nanoESI (+) and a NanoLock Spray (Waters Corporation, Manchester, UK) ionization source. The lock mass channel was sampled every 30 s. Glu1-Fibrinopeptide B human solution was used to calibrate the mass spectrometer with an MS/MS spectrum reference using the NanoLock Spray source. Samples were all run in biological replicates.

### Data processing and quantification

High-definition data-independent MS raw files were processed for label-free identification and quantification in Progenesis® QI for proteomics version 4.0, including software Apex3D, Peptide 3D, and ion accounting informatics programs (Waters). Starting with LC-MS dataset loading, the software performed alignment and peak detection, providing a list of peptide ions (peptides) to be explored with multivariate statistical methods in Peptide Ion Stats. Finally, proteins were identified with dedicated algorithms and cross-matching with the Uniprot human proteome database (version 2018/09); default parameters for ion accounting and relative quantitation with the Hi-N (3) method of peptide comparison were used ^40, 41^. To remove false-positives, reversed database queries were appended to the original database. The protein/peptide level false discovery rate was set at 1%. Digestion by trypsin allowed a limit of one missed cleavage. Methionine oxidations and carbamido-methylations were considered variable and fixed modifications, respectively. Identifications that did not satisfy these criteria were rejected.

### In silico analyses

Differentially regulated proteins were subjected to functional and enrichment analyses in Metascape ^42^, which searches databases such as Reactome Pathways Knowledgebase (https://reactome.org; ^43^) and the KEGG knowledge base (kegg.jp; ^44^). Protein interaction networks were identified with the help of STRING database (https://string-db.org; ^45^). Visual integration was performed in Cytoscape v3.8.2 (https://cytoscape.org; ^46^). Ingenuity Pathway Analysis (Qiagen Bioinformatics, Redwood, CA) was used to perform a core analysis of diseases and biological functions. Principal component analysis was performed on log-transformed data, enabling multivariate comparisons among groups in ClustVis (biit.cs.ut.ee/clustvis), which was employed with default parameter settings ^47^. Hierarchical clustering and enrichment of secretome data were performed in ClusterExp (https://fgvis.com), which searches the KEGG and Reactome databases.

### CAM assay

For *in vivo* evaluation of the angiogenic inductive potential of astrocyte secretomes, CAM assays were performed ^38^. Briefly, fertilized chicken eggs (Agricola Chorombo, Chile) were incubated at 38.5 °C under constant humidity. At embryonic day 1 (E1), 3 ml of albumin was extracted from each egg; a round window (2 cm^2^) was created on E4. A bio-cellulose scaffold 6 mm in diameter was filled with 100 μl of medium to be assayed: conditioned medium from astrocytes, DMEM/F12 (negative control), and 100 μg VEGFA (positive control). Human recombinant IL-8 (catalog no. 208-IL-010, R&D Systems, USA) was used at a 5 ng/ml concentration. On E8, the CAM vasculature was photographed; subsequently, each experimental condition scaffold was placed on top of the CAM. For each condition, 10 eggs were used. On E12, white cream was injected under the CAM before photographing to improve vessel visualization. Photographs were taken with a digital camera (Leica HD IC80, Heidelberg, Germany). To identify and quantify the changes generated in the CAM vasculature, vessel number and thickness within a 6-mm radius of the scaffold were quantitated. Vessel morphology was examined in ImageJ software (NIH, USA).

### Statistical analyses

Comparisons between two groups were made with unpaired Student’s t‐tests. Comparisons between more than two groups were made with one-way or two-way analyses of variance (ANOVAs; treatment and disease as factors), followed by Tukey’s post hoc tests. The criterion for significance was p < 0.05. Data analysis was performed in Prism v8.02 (GraphPad Software, La Jolla, CA). Proteomics data were subjected to one-way ANOVAs (p ≤ 0.05 as considered differentially regulated) with biological replicate and experimental condition treated as independent factors.

## Results

### Immunocytochemistry and proteomic analyses of hiPSC-derived astrocytes

Morphology and astroglial marker presence were examined in control and SCZ hiPSC-derived astrocytes with immunocytochemistry (Figure 2A) and proteomic analysis (Figure 2B-E). Differences between control and SCZ hiPSC-derived astrocytes were detected via a discovery-driven proteomics approach. Label-free proteomic analyses identified a total of 1,868 proteins, of which 1,636 were present in both SCZ and control groups (Figure 2B and Supplementary Table 2). Of these, 68 proteins were found to be significantly dysregulated in the SCZ group (p < 0.05 vs. control) (Figure 2B). Cells from both groups were GFAP- and vimentin-immunopositive (Figure 2A & C), confirming they contained astrocyte-typical intermediate filaments. Similar abundances were observed for membrane markers, including the glial glycoprotein CD44 and ITGA6 (integrin alpha 6), a putative astrocyte cell-surface marker ^48^ (Figure 2C). Both groups of cells were also confirmed to be expressing excitatory amino acid transporters 1 and 2 (EAAT1 and EAAT2) (Figure 2A) as well as Na^+^/K^+^ ATPase (ATP1A2) (Figure 2C), proteins that are critical for the astroglial modulation of glutamate/aspartate neurotransmission. We also found positive labeling for Connexin 43 (Cx43) (Figure 2A), an important gap junction protein highly expressed by astrocytes. Proteins related to astroglial metabolism, such as mesencephalic astrocyte-derived neurotrophic factor (MANF), which is secreted by astrocytes, and the glycolytic enzyme ALDOC (a class I fructose-bisphosphate aldolase specific to astrocytes) were similarly expressed in both groups (Figure 2C). Lastly, control and SCZ hiPSC-derived astrocytes presented labeling for aldehyde dehydrogenase-1 family member L1 (ALDH1L1), an antigenic marker specific for astrocytes and astroglial progenitors within the CNS (Figure 2A). All tested astroglial markers were abundantly present in cell lines with 98% of cell positivity in every cell line tested (data not shown).

**Figure 1.**
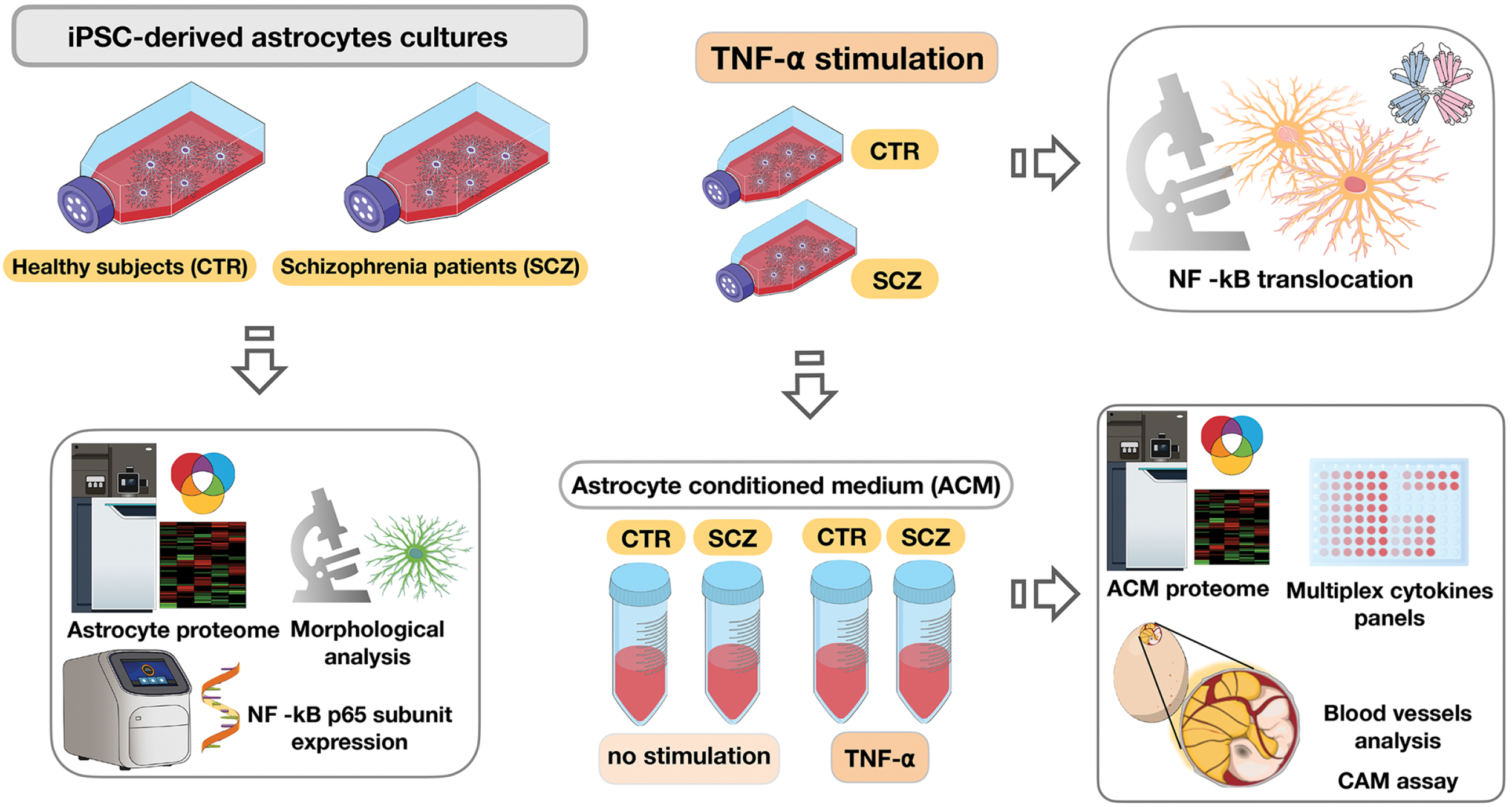
Workflow sketch. Human induced pluripotent stem cells (hiPSC)-derived astrocytes obtained from patients diagnosed with schizophrenia (SCZ) and control (CTR) subjects were submitted to shotgun proteomic analysis. In order to confirm the biological identity of these cells, and if schizophrenia would affect them, hiPSC-derived astrocytes from both groups were immunolabeled for classical astrocyte markers in which part of these targets were previously detected in the proteomics. Following alterations in pathways related to inflammation pointed by systems biology analysis, NF-kB p65 subunit expression was analyzed. Subsequently, NF-kB p65 subunit nuclear translocation was evaluated in CTR- and SCZ-hiPSC-derived astrocytes stimulated with TNF-α. Astrocyte conditioned media (ACM) from unstimulated and TNF-α-stimulated hiPSC-derived astrocytes from both CTR and SCZ groups were further analyzed for inflammation-related cytokines and also submitted to shotgun proteomics. Finally, we tested the hypothesis that angiogenesis might be affected by ACM from SCZ patients by testing, *in vivo*, ACM from all experimental groups through the Chicken chorioallantoic membrane (CAM) assay.

**Figure 2.**
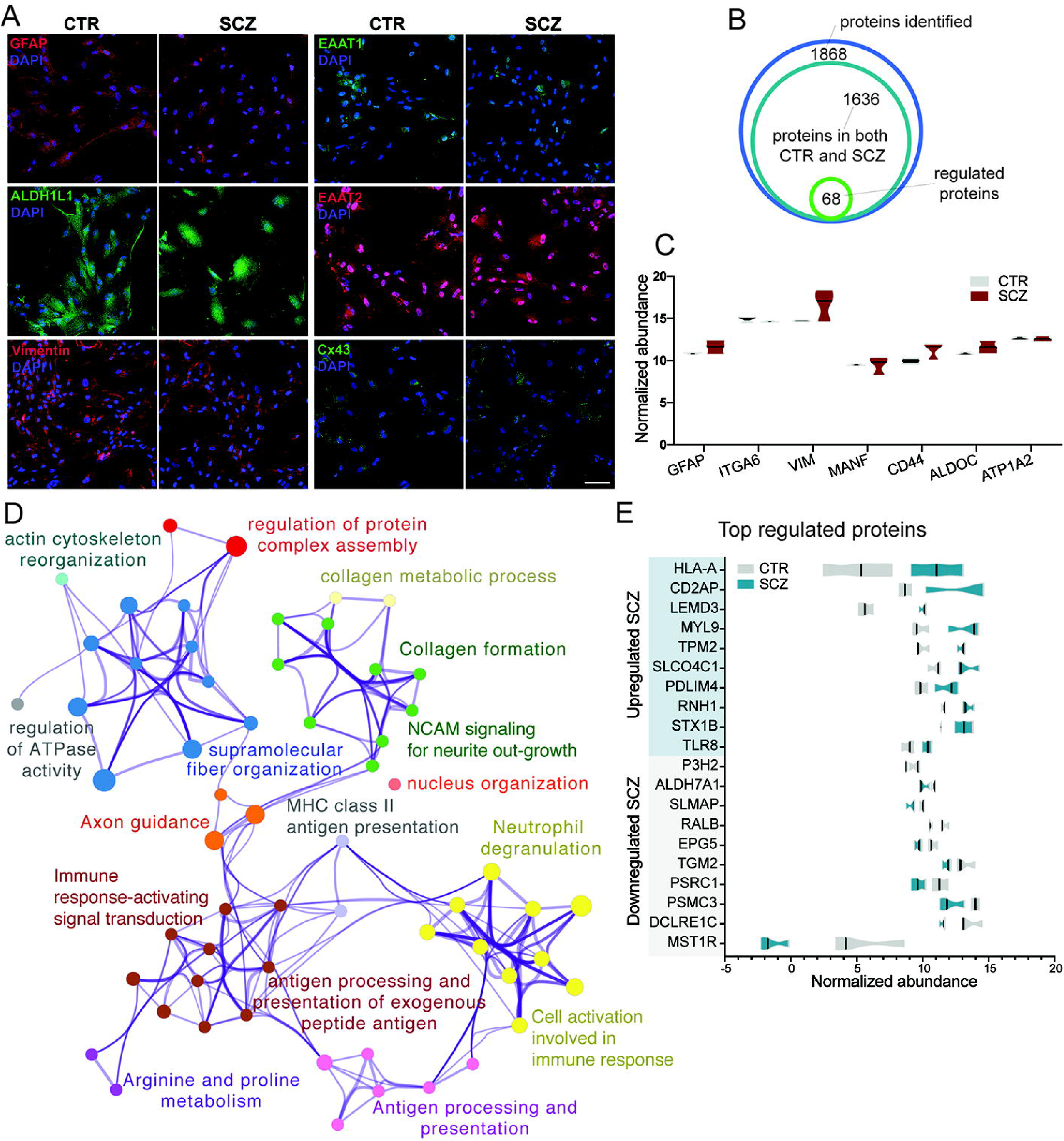
hiPSC-derived SCZ astrocytes exhibit similar protein markers as compared to CTR and differential regulation of pathways. (A) Immunocytochemistry for astrocyte markers (GFAP, ALDH1L1, Vimentin, EAAT1, EAAT2, Cx43). Astrocyte markers were found in most hiPSC-derived astrocytes. No differences were observed between CTR and SCZ groups. (B) Venn diagram of the number of proteins identified and quantified by mass spectrometry-based shotgun proteomics. The diagram shows the regulated proteins in hiPSC-derived SCZ astrocytes in comparison to CTR. (C) Proteomics quantification of astroglial protein markers. The comparison (Log_2_) was a normalized abundance of CTR and SCZ astrocytes, and no significant differences were observed between them. (D) Network representation of statistically enriched clusters (including GO terms, Reactome, and KEGG pathways). Each node represents a significant term, which is colored by hierarchical cluster-ID. Nodes that share the same cluster are connected and colored alike. The top term was used as cluster representation. The network was built using Metascape (https://metascape.org) and modified with Cytoscape (v3.1.2). (E) Proteomics quantification of the ten top significantly regulated proteins (up- and downregulated) in SCZ and CTR astrocytes. Several of those proteins were clustered to inflammation-associated pathways. The normalized abundance of proteins (Log_2_) is shown, only different proteins (p< 0.05) are shown. Magnification of photomicrographs: 200x. Calibration bar: 100 μm.

Differentially regulated proteins previously revealed in proteomics studies that compared SCZ and control hiPSC-derived astrocytes have been related to ontological groups and pathways related to cell reorganization and have been found in studies of SCZ brain cells (for review, see ^27^). These include proteins related to cytoskeletal actin, regulation of protein complex assembly, ATPase activity, collagen formation, and metabolic processes (Figure 2D). Additionally, some of the differentially regulated proteins have been related to astrocyte regulation of CNS development, including axon guidance and immune-response cell activation.

Several proteins associated with the innate immune response were found to be upregulated in SCZ astrocytes, compared to controls, including HLA-A (HLA class I histocompatibility antigen, A-32 alpha chain) and toll-like receptor 8 (TLR8), as well as syntaxin-1B (STX1B), which is involved in BDNF release (Figure 2E). Some proteins that regulate protein degradation, migration, and proliferation were found to be down-regulated, such as the 26S proteasome regulatory subunit 6A (PSMC3) and macrophage-stimulating protein receptor (MST1R) (Figure 2E). Importantly, these dysregulated proteins play essential roles in cell development and participate in glial support of CNS homeostasis. Proteins involved in antigen presentation pathways, neutrophil degranulation, and immune response-activating signal transduction biological processes were also found to be dysregulated in SCZ astrocytes (Figure 2D).

### NF-κB p65 subunit is down-regulated in SCZ hiPSC-derived astrocytes

Analysis of protein-protein interaction networks related to innate immune signaling revealed up- and down-regulated proteins interconnected within networks. Further analysis of these interactions revealed markers related to inflammation, including S100β and the transcription factor NF-κB (Figure 3A). These findings prompted us to conduct experiments designed to examine the response of SCZ-derived astrocytes to inflammation and NF-κB p65 expression and nuclear translocation, which is important for eliciting cytokine production.

**Figure 3.**
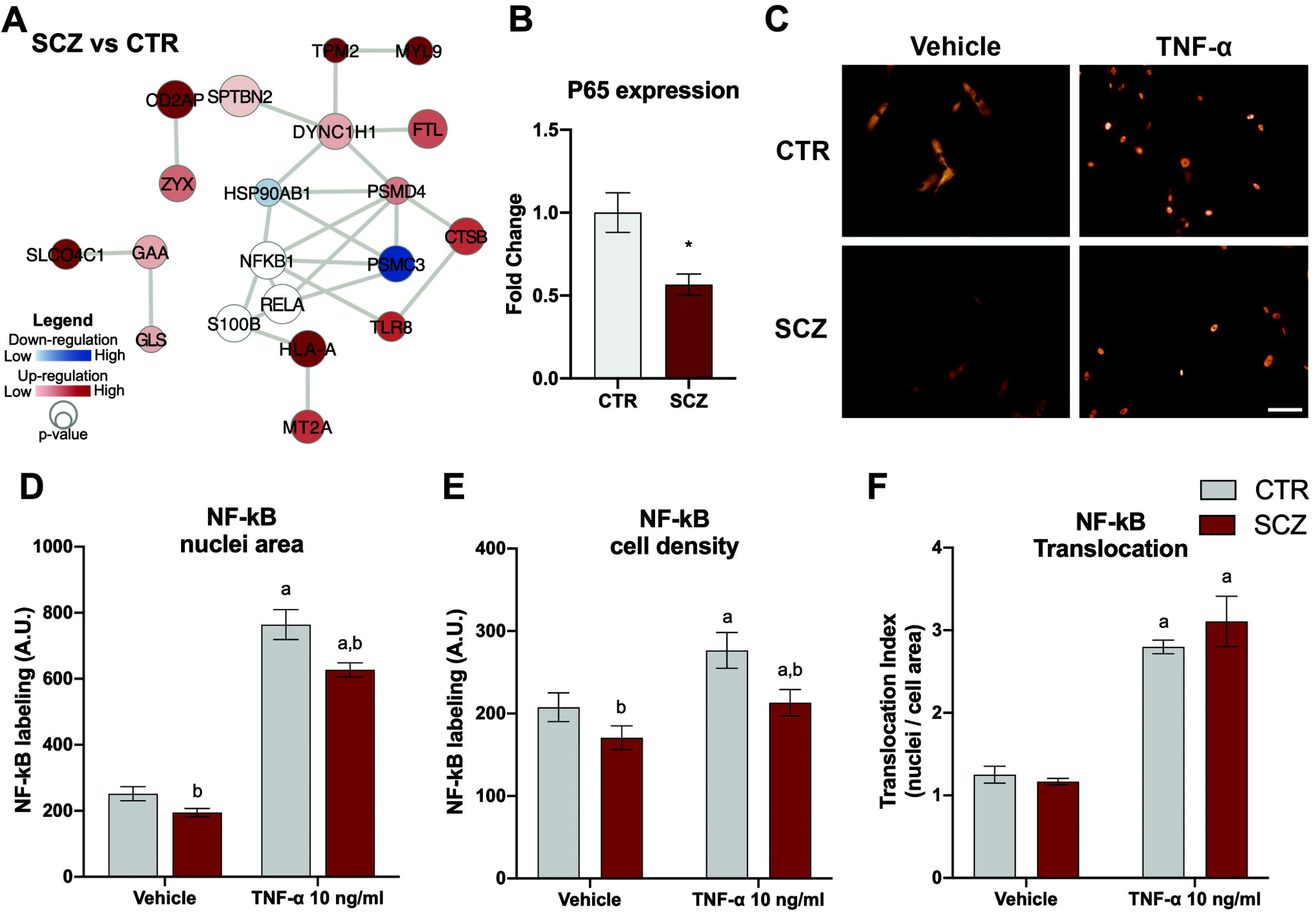
hiPSC-derived SCZ astrocytes inflammatory phenotype and p65 subunit of NF-κB response to TNF-α. (A) Protein-protein interaction network of upregulated (colored in red) and downregulated (colored in blue) inflammation-related proteins in SCZ in comparison to CTR astrocytes. Proteins regulated were connected to NF-κB (RELA/p65 and NFKB1/p50) and S100B (uncolored, not found in the proteome dataset). Interaction networks are based on the STRING database (https://string-db.org) and Reactome (https://reactome.org). (B) Real-time PCR for NF-κB p65 subunit expression in cell extracts from hiPSC-derived astrocytes. (C) Photomicrographs of NF-κB p65 subunit immunostaining 1 h after exposing cells to vehicle or 10 ng/ml TNF-α. Quantification of NF-κB p65 subunit immunoreactivity in (D) nuclei and (E) cell area; and (F) NF-κB translocation index (nuclei/whole cell area ratio). Magnification of photomicrographs: 100x. Calibration bar: 100 μm. Experiments were performed in triplicates from 3 cell lines per condition. Data in (B) are presented as means ± SEM; *p<0.05, Unpaired Student’s t-test. Data in (D, E, F) are presented as means ± SEM; Two-way ANOVA, followed by Tukey’s multiple comparisons test: a – treatment effect (p<0.01), TNF-α different from vehicle; b – disease effect (p<0.01), difference between CTR and SCZ cells.

RT-qPCR experiments conducted to evaluate NF-κB responses following TNF-α stimulation showed that NF-κB p65 expression was decreased by ~50% in SCZ hiPSC-derived astrocytes compared to control astrocytes (Figure 3B), corroborating an effect on NF-κB regulation predicted by the protein network interaction analysis. Both control and SCZ astrocytes showed nuclear translocation of NF-κB following 1-h TNF-α (10 ng/ml) exposure (Figure 3C & F). Densitometric analysis showed a 20% reduction of NF-κB immunolabeling in both nuclei and cell areas of SCZ hiPSC-derived astrocytes (Figures 3D & E).

### Angiogenesis-related proteins are disrupted in SCZ hiPSC-derived astrocytes

The reduction of NF-κB p65 subunit in SCZ hiPSC-derived astrocytes led us to test whether inflammation responses could be disrupted in these cells. Using MS-based shotgun proteomics, we compared A_SCZ_CM (i.e., SCZ astrocyte secretome) between resting state (no stimulation) and TNF-α stimulation conditions, with the latter being expected to trigger NF-κB translocation. In total, we identified 1,064 proteins, 863 of which remained following application of the exclusion criteria (e.g., being found in both non-stimulated and stimulated cells). Of those, 64 proteins were found to be significantly differentially regulated (p < 0.05) among the experimental groups (Figure 4A and Supplementary Table 3).

**Figure 4.**
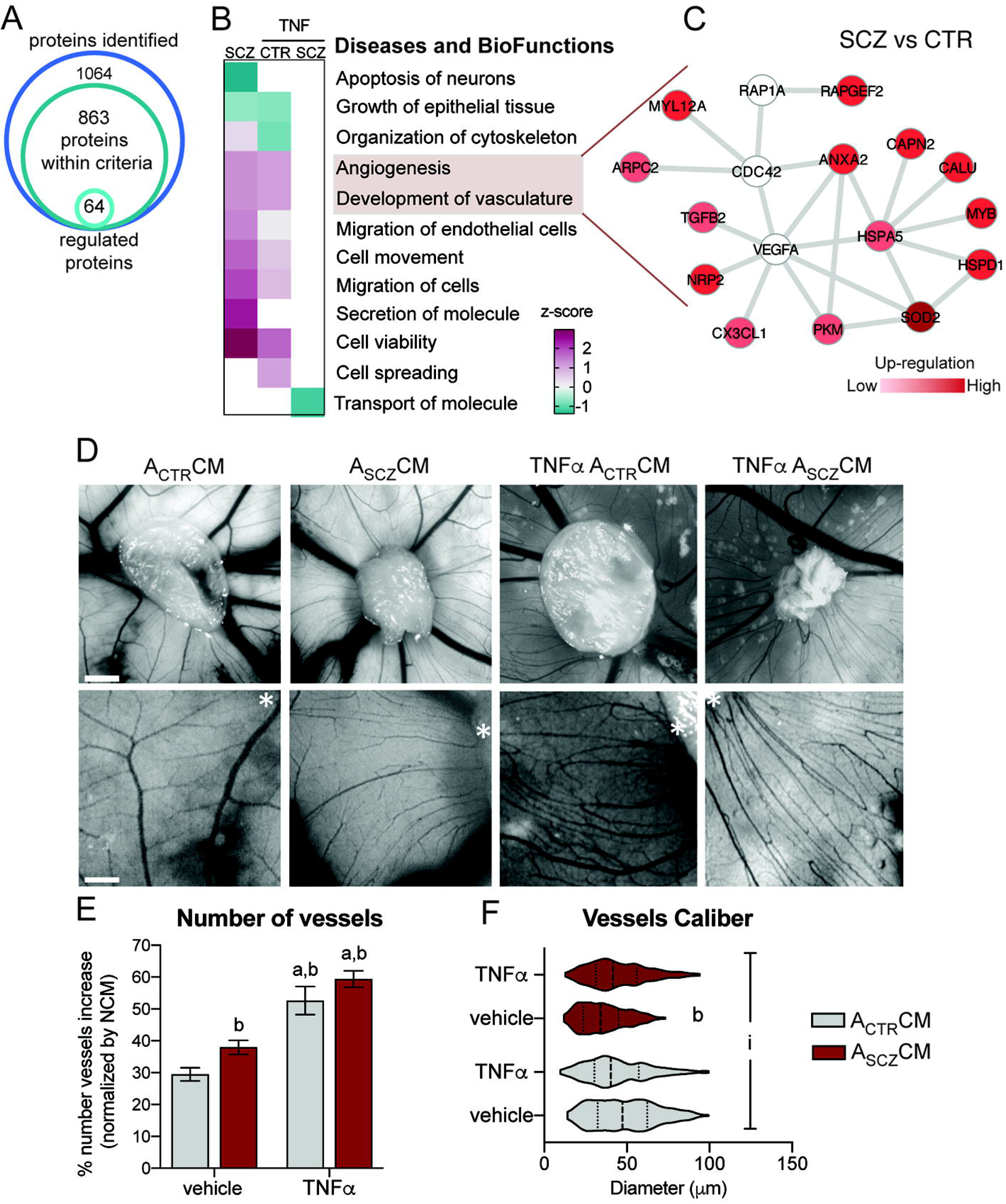
The secretome of hiPSC-derived SCZ astrocytes and its effect on blood vessel development in the chicken chorioallantoic membrane. (A) Venn diagram showing the number of proteins identified and quantified by mass spectrometry-based shotgun proteomics in astrocytes conditioned medium. The total proteins of regulated A_SCZ_CM and A_CTR_CM treated with vehicle and TNF-α are represented. (B) Enrichment heatmap of diseases and bio-functions terms and activation for significantly regulated (p<0.05) A_SCZ_CM versus A_CTR_CM; TNF-α-treated A_SCZ_CM and A_CTR_CM versus vehicle ACM. Colors indicate z-score, red for activation and green for inhibition. Functional activation calculated on Ingenuity Pathway Analysis ®. (C) Interaction network of proteins in A_SCZ_CM versus A_CTR_CM related to enrichment analysis of the angiogenesis and development of vasculature bio-functions. The regulated proteins interact-partners include VEGFA, CDC23, and RAP1A (uncolored, not found in the proteomics dataset). Protein nodes upregulated (p<0.05) are colored in red. Interaction networks are based on the STRING database (https://string-db.org) and modified on Cytoscape (v3.1.2). (D) Representative images of CAM vessels when the scaffold was loaded with A_CTR_CM (vehicle and TNF-α-treated), and A_SCZ_CM (vehicle and TNF-α-treated). Asterisks (*) in bottom images indicate a reference to the scaffold position which is an important landmark to the vessels’ analyses. (E) Quantification of the increase in the number of vessels in each condition at E12 compared to E8 (normalized to control). (F) Violin plot exhibiting the distribution of the vessel’s caliber found in the different experimental conditions. Vertical bars represent the mean vessel caliber for each experimental group. CTR and SCZ hiPSC-derived astrocytes were stimulated for 24 h with 10 ng/ml of TNF-α or vehicle. Conditioned medium was collected from four CTR cell lines and three SCZ cell lines, each individually assayed. Upper calibration bar: 2 mm. Bottom calibration bar: 70 μm. Data are presented as mean ± SEM; Two-way ANOVA, followed by Tukey’s multiple comparisons test; a – treatment effect (p<0.01), TNF-α different from vehicle; b – disease effect (p<0.05), difference between CTR and SCZ cells; i - interaction between (p<0.001), treatment and disease (p<0.001).

Analyzing relative fold changes between SCZ versus control groups and TNF-α stimulated versus non-stimulated cells revealed similar modulation across the non-stimulated SCZ cells and the TNF-α stimulated control cells (Figure 4B). The biological functions implicated include inhibition of neuronal apoptosis and epithelial tissue growth, activation of angiogenesis, development of vasculature, cell migration, and cell viability (Figure 4B). Some proteins found to be regulated in the secretome of SCZ cells stimulated with TNF-α are related to inhibition of molecule transport (Figure 4B). Proteins involved in the angiogenesis and development of vasculature were upregulated in non-stimulated SCZ hiPSC-derived astrocytes (vs. control, no stimulus), including several in a neighboring network of VEGFA, including the TGF-α family protein TGFB2, annexin (ANXA2), neuropilin (NRP2), and fractalkine (CX3CL1) (Figure 4C).

Because astrocytes exert tight BBB control, in part through strict regulation of angiogenesis, we tested whether molecules present in A_SCZ_CM and A_CON_CM would modulate angiogenesis in the chorioallantoic membrane (CAM) *in vivo* assay. Angiogenic properties were examined between E8 and E12, a critical period of vessel development during which robust changes in CAM vessels can be observed. The CAM assay analysis demonstrated that conditioned media induced significantly more angiogenesis than vehicle, with the A_SCZ_CM treatment resulting in significantly more vessels than A_CON_CM treatment within both resting and TNF-α-exposed conditions (Figures 4D & E). TNF-α altered vascular morphology, including changes in the vessel network, in both A_SCZ_CM and A_CON_CM assays (Figure 4D). Compared to A_CON_CM, A_SCZ_CM reduced the mean CAM vessel caliber by ~30%; TNF-α had similar effects on caliber in both groups (Figure 4F).

### Inflammation-related cytokine secretion is disturbed in SCZ hiPSC-derived astrocytes

Principal component analysis did not reveal any clear clusters when comparing four groups composed of control and SCZ hiPSC-derived astrocytes under resting state (no stimulation) and activated by TNF-α (Figure 5A). While the control (no stimulation) group showed some separation from the other experimental conditions, the non-stimulated control and SCZ secretomes exhibited the most differentiated regulation of the 64 significantly regulated proteins. Several proteins that are upregulated in SCZ cells, compared to control cells, showed no major differences in response to TNF-α exposure (Figure 5B). Hierarchical clustering of those proteins showed that TNF-α stimulation upregulated cellular processes such as mitosis and vesicle-mediated transport independent of group. Independent of TNF-α stimulation, SCZ hiPSC-derived astrocytes showed upregulation of proteins related to hypoxia response, MAPK signaling, the VEGF-VEGFR2 pathway, nerve growth factor response, cAMP, and regulation of cytokine production, among others (Figure 5B).

**Figure 5.**
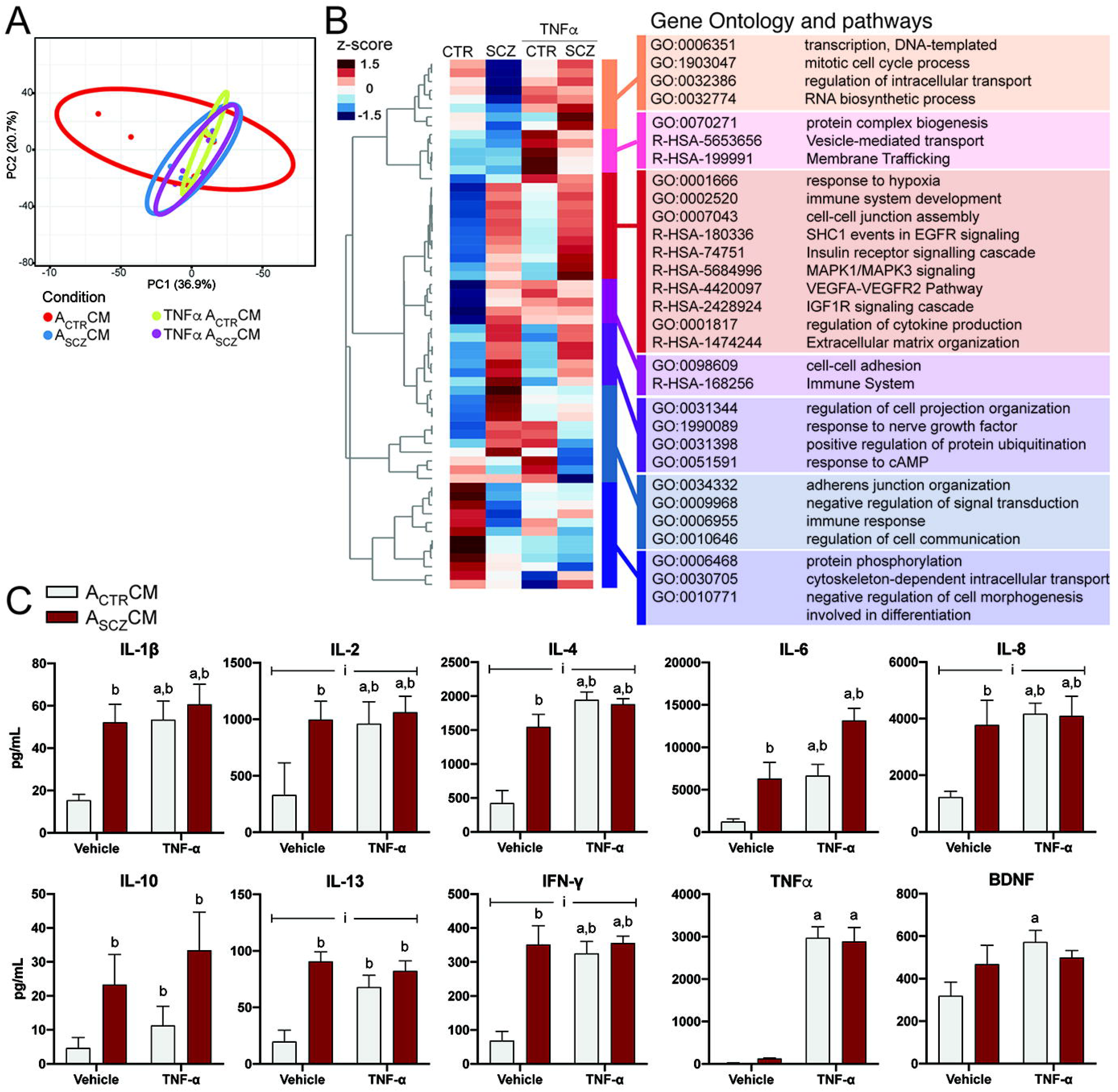
Regulation of the secretome of hiPSC-derived astrocytes reveal clusters of inflammation-related proteins and increased levels of inflammatory cytokines. (A) Principal component analysis of the protein regulation in A_SCZ_CM and A_CTR_CM exposed to TNF-α or vehicle. Shapes are representing each sample and ellipses clustering of those samples. Clustering shows a separation of A_CTR_CM, while A_SCZ_CM and TNF-α-treated A_SCZ_CM and A_CTR_CM cluster together. (B) Heatmap with the hierarchical clustering of differentially regulated proteins (p<0.05) in A_SCZ_CM and A_CTR_CM exposed to TNF-α or vehicle. Protein downregulation is represented in blue and upregulation in red. Cluster groups of proteins were then selected to search for enriched gene ontology and pathways (Reactome, KEGG, GO terms were considered), and are grouped by different colors. (C) Cytokine secretion levels in the A_CTR_CM and A_SCZ_CM resting cells (vehicle) and TNF-α-exposed cells. Multiplex measurement of Interleukin-1 beta (IL-1β), Interleukin-2 (IL-2), Interleukin-4 (IL-4), Interleukin-6 (IL-6), Interleukin-8 (IL-8), Interleukin-10 (IL-10), Interleukin-13 (IL-13), Interferon gamma (IFN-γ), Tumor necrosis factor alpha (TNF-α), and brain-derived neurotrophic factor (BDNF). Data show the concentration (pg/mL) of cytokines in astrocyte conditioned media. CTR and SCZ hiPSC-derived astrocytes were stimulated for 24 h with 10 ng/ml of TNF-α or vehicle. Conditioned media were collected from four CTR cell lines and three SCZ cell lines. Experiments were performed in duplicates. Data are presented as means ± SEM. Two-way ANOVA followed by Tukey’s multiple comparisons test was used; a – treatment effect (p<0.01), TNF-α different from vehicle; b – disease effect (p<0.05), difference between CTR and SCZ cells; i – interaction between, treatment and disease (p<0.05).

To determine which secreted cytokines may underlie the above effects, we performed multiplex analysis of released cytokines in the conditioned media under resting and TNF-α stimulated states, as in the above experiment. We found that, compared to A_CON_CM, the A_SCZ_CM had pronounced and significant increases in all inflammation-related cytokines (Figure 5C), consistent with a proinflammatory effect. Significant differences in BDNF between the A_CON_CM and A_SCZ_CM groups were detected only after TNF-α stimulation. Interestingly, TNF-α stimulation increased the expression of several cytokines in A_CON_CM, while levels observed in A_SCZ_CM were similar to levels seen in non-stimulated cells (Figure 5C). Consequently, a significant interaction effect was shown between disease and stimulation for the cytokines IL-2, IL-4, IL-8, IL-13, and IFN-L. Notably, IL-8 was found at particularly high levels in A_SCZ_CM. As expected, IL-6 was highly induced in A_SCZ_CM, in which it was present at 66% higher levels than in the A_CON_CM following TNF-α activation.

### IL-8 mimics the vascularization disruption of SCZ hiPSC-derived astrocytes in chicken embryos

Several inflammation-related proteins were differentially regulated in the secretome of SCZ iPSC-derived astrocytes and this secretome influenced angiogenesis in CAM assays. Because the chemokine IL-8 has been associated with chemotaxis and modulation of endothelial function ^49^ was found in high concentrations in non-stimulated A_SCZ_CM (Figure 5C), we tested whether human recombinant IL-8 alone could reproduce the angiogenesis changes induced by A_SCZ_CM (Figure 6A). Exogenous addition IL-8 (5 ng/ml) increased the number of vessels to an extent similar to that seen with A_CON_CM (Figure 6B). When IL-8 was provided together with A_CON_CM, the number of vessels observed was similar to the quantities recorded with A_SCZ_CM (Figure 6A & B). Likewise, exogenous IL-8 reduced mean vessel caliber to an extent similar to that observed with A_SCZ_CM (Figure 6C). The A_SCZ_CM and exogenous IL-8 conditions yielded similar vessel caliber distributions (see violin plots in Figure 6C), suggesting that IL-8 may be a major regulator of vessel caliber for the A_SCZ_CM condition.

**Figure 6.**
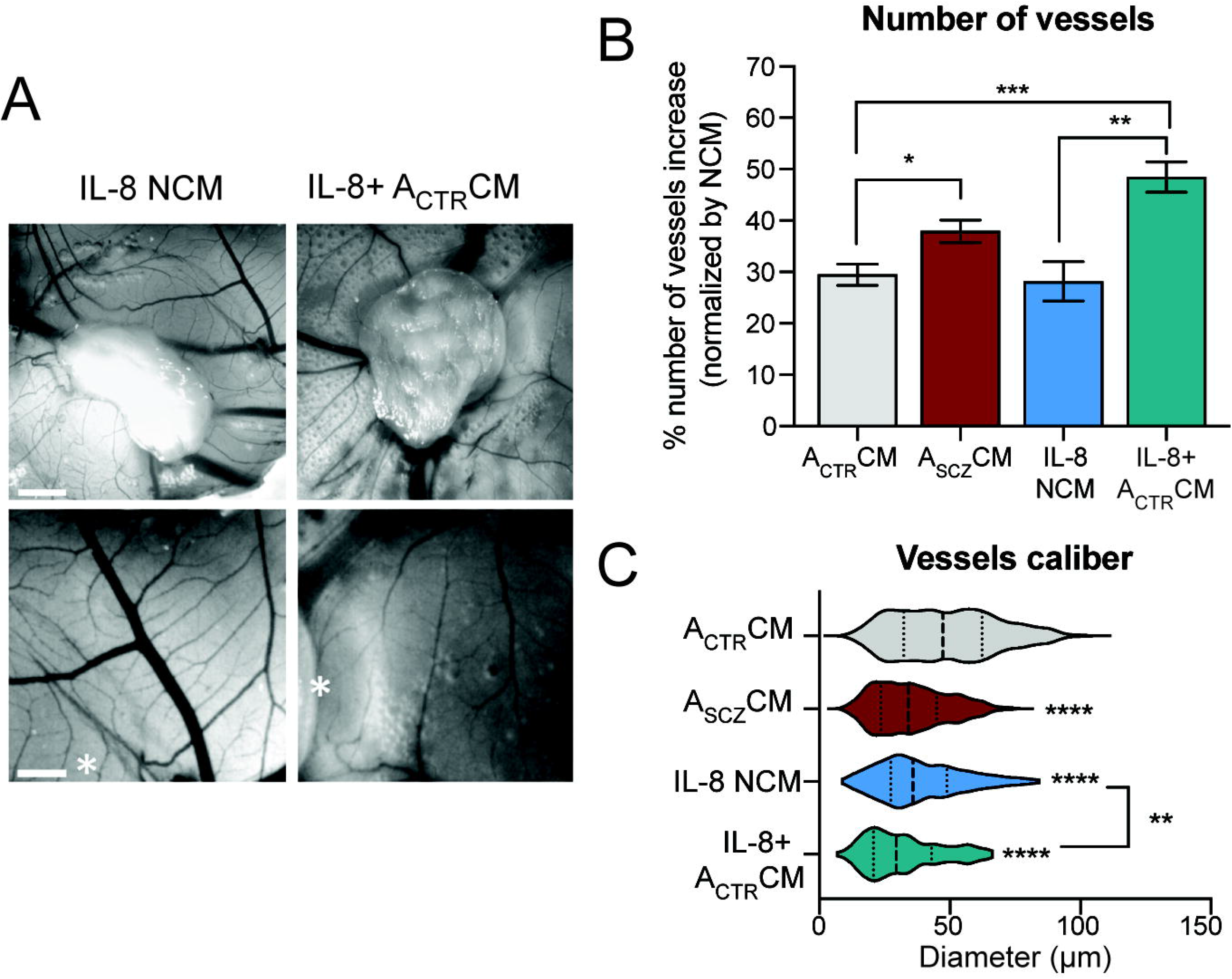
Conditioned Media from hiPSC-derived SCZ astrocytes and Interleukin-8 increased the number and reduced the caliber of vessels in the chicken chorioallantoic membrane. (A) Representative images of CAM vessels when the scaffold was loaded with vehicle (DMEM/F12) + 5 ug/ml IL-8 or A_CTR_CM + 4 ug/ml IL-8, as indicated. Asterisks (*) in bottom images indicate a reference to the scaffold position which is an important landmark to the vessels’ analyses. Upper calibration bar: 2 mm. Bottom calibration bar: 70 μm. (B) Quantification of the increase in the number of vessels in each condition at E12 compared to E8 (normalized to control). A_CTR_CM and A_SCZ_CM (Fig. 4) are included for comparison. (C) Violin plot exhibiting the distribution of the vessel’s caliber found in the different experimental conditions. Vertical bars represent the mean vessel caliber for each experimental group. Conditioned media were collected from four CTR cell lines and three SCZ cell lines. CAM assay was performed in a minimum of six eggs per experimental group. Two hundred vessels were analyzed in each group. *p<0.05; **p<0.01; ***p<0.001. One-way ANOVA followed by Tukey’s multiple comparisons test was used.

## Discussion

Here we showed that hiPSC-derived astrocytes obtained from patients diagnosed with SCZ exhibited a chronic inflammatory profile with broad effects on its secretome. There is a large body of evidence suggesting that neuroinflammation may be a major player in the etiology of SCZ (for review, see ^50^). Both our proteomic and cytokine array analyses revealed alterations in pathways of vascularization signaling, highlighting the chemokine IL-8 as a major candidate in SCZ-associated vascular dysfunction. We found alterations in immune system-related proteins in SCZ hiPSC-derived astrocytes, including alterations in TLR8 and human leukocyte antigen (HLA)-A expression, corroborating previous observations of enhanced TLR8 activity in peripheral blood cells from patients with SCZ ^51^. Meanwhile, genome-wide association studies have revealed an association of SCZ with polymorphisms or changes in HLA expression ^52–54^.

We did not detect significant control subject versus SCZ patient differences in the astrocyte markers ALDOC and GFAP, although these proteins have been reported to be affected in SCZ in postmortem studies ^26, 27^. Given that our study was based on hiPSC-derived astrocytes, rather than postmortem brains, it may be that the alterations observed in postmortem SCZ brains reflect more complex interactions of astrocytes with other cell types.

In the interactome map, both NFKB1 and RELA (p65) are predicted targets of immune-related proteins likely to be affected in SCZ astrocytes, thus representing a pivotal hub in cascade signaling. This outcome fits with prior observations of reduced NF-κB p65 subunit levels in SCZ astrocytes as well as with NF-κB pathway abnormalities that have been found in postmortem SCZ brain samples ^55^. Although our SCZ astrocytes presented constitutively reduced NF-κB p65, increased amounts of inflammatory and modulatory cytokines were present in A_SCZ_CM and nuclear translocation of NF-κB p65 was observed after TNF-α exposure. NF-κB1 precursor can be processed into the mature p50 subunit, which partners with p65 to form the major heterodimer in the canonical NF-κB activation pathway. Of the several NF-κB dimer conformations that can form among NF-κB subunits, including homodimers (Murphy et al., 2021), the p50/p65 heterodimer is thought to be the predominant activated form of NF-κB in glia ^56^. There remains a need for more studies of NF-κB dimer compositions in SCZ astrocytes.

Cytokine imbalances have been reported frequently in SCZ ^57^. Despite the low expression of NF-κB p65, we found that A_SCZ_CM had increased amounts of inflammatory and modulatory cytokines, similar to that of TNF-α stimulated A_CON_CM. With the exception of IL-6, SCZ hiPSC-derived astrocytes stimulated with TNF-α did not show any incremental increase in cytokines, which may lead these cells to adopt an activated status wherein they are unable to respond to TNF-α. This chronic immune activation exhibited by SCZ hiPSC-derived astrocytes may also affect the NF-κB pathway in these cells through negative feedback regulation. These results corroborate the findings of Akkouh and collaborators (2020), who showed that SCZ hiPSC-derived astrocytes exhibited an impaired CCL20 response to IL-1β stimulation, which might affect the migration of regulatory T cells and brain inflammation.

Our analyses of astrocyte conditioned media showed that TNF-α induced a modest effect on the modulation of several pathways and functions related to immune regulation in SCZ hiPSC-derived astrocytes. Yet, principal component analysis of secretomes revealed that SCZ hiPSC-derived astrocytes, stimulated or not with TNF-α, clustered very closely with control subject hiPSC-derived astrocytes exposed to TNF-α. Previously, we showed that conditioned media of hiPSC-derived NSCs from SCZ subjects exhibited disrupted secretion and expression of angiogenic factors ^38^. Windrem and collaborators showed that SCZ hiPSC-derived glial progenitors exhibited disrupted gene expression of targets related to synaptic regulation and glial differentiation. In addition, they found that SCZ hiPSC-derived glial progenitors exhibited delayed astrocytic differentiation when xeno-transplanted into the corpus callosum of mice, which was associated with behavioral and electrophysiological effects *in vivo* ^36^. Despite the technical distinctions between differentiation protocols, the hiPSC-derived cells used in Windrem’s and Casas’ works are both glial progenitors, suggesting that the glial cell lineage may be intrinsically disrupted during brain development in SCZ. Collectively, these data suggest SCZ-associated dysregulation of neuroinflammatory pathways, often observed in adult brains ^58^ and intrinsically related to gene regulation in astrocytes ^59^, would have some origins apparent during development ^60^.

Secreted proteins may provide clues regarding how dysregulation of SCZ astrocytes may affect the cellular microenvironment. For instance, simultaneous upregulation of proteins such as TGFB2, NRP2 (glycoprotein neuropilin), PKM (pyruvate kinase 2), and SOD2 (superoxide dismutase) suggest that cellular movement, vasculogenesis, and angiogenesis may be affected. Of note, several of these proteins are involved in vascular and neural development ^61, 62^. Neuropilin acts as a co-receptor in both pathways during axon morphogenesis ^63^ and angiogenesis (Gelfand et al., 2014), as well as in association with semaphorin and VEGF. Here, we observed that all experimental groups of hiPSC-derived astrocyte-conditioned media increase angiogenesis in chick embryos, supporting the supposition that astrocytes are an important source of angiogenic factors in the CNS ^64^. Our finding of higher levels of angiogenesis and decreased vessel caliber with A_SCZ_CM than with A_CON_CM suggests astrocytes may play a major role in the disruption of vascularization in SCZ. On the other hand, we previously showed that conditioned media from SCZ NSCs exhibited weak angiogenesis induction ^38^. Compared to astrocytes, NSCs are expected to have a slightly inflammatory response. Moreover, adult NSCs have latent inflammatory potential that is chronically suppressed to facilitate neurogenesis ^65^. Therefore, we suggest that the increased angiogenesis induced by SCZ astrocytes may be a byproduct of inflammatory disruption.

Vascular alterations are common features of SCZ ^66, 67^. The authors of neuroimaging studies have described microvasculature abnormalities in SCZ, including reduced capillary area density in the prefrontal cortex, but not in the visual cortex ^68^. In the retina, wider venules were reported in SCZ ^69, 70^. These vascular changes correlate with reduced cortical thickness in patients diagnosed with SCZ ^71^. Modulation of vessel caliber might impact local neuronal circuitry by affecting blood flow. SCZ has been reported to be associated with both up- or down-regulation of blood flow in the thalamus, cerebellum, striatum, and cortex ^72–75^. Observations of wide and differentially distributed CNS vascular alterations in SCZ underscore the need for further investigations into the physiological aspects of distinct astrocyte populations across these regions.

The chemokine IL-8, which we found to be highly upregulated in SCZ astrocytes, was sufficient to increase vessel formation in a manner similar to A_SCZ_CM, including the smaller caliber vessels commonly associated with angiogenesis ^49^. Elevated IL-8 levels during pregnancy have been associated with an increased risk of SCZ in the adult offspring ^20, 76, 77^. High levels of IL-8, as well as IL-6 and IL-2, have also been observed in the plasma of untreated SCZ patients ^78–80^. This chemokine is a potent chemoattractant for lymphocytes and neutrophils during inflammation ^81^, and may act as a key player in multiple brain conditions, including Alzheimer disease ^82^ and depression ^83^. IL-8 was shown to affect pericytes, allowing them to chemoattract neutrophils when activated by inflammatory stimuli ^84^. Intra-cerebrospinal fluid injection of IL-8 is associated with BBB dysfunction following brain injury ^85^. Moreover, IL-8 has been identified as a trophic factor that promotes endothelial cell survival and proliferation as well as a pro-angiogenic factor ^86, 87^. Therefore, it is possible that IL-8 could be a central player in neuroinflammation and vascular alterations in SCZ by way of its effects on the BBB, which might give circulating immunocompetent cells access to the CNS. Previous findings of an elevated neutrophil count ^88, 89^ and an elevated neutrophil-to-lymphocyte ratio ^90^ in the peripheral blood of patients with SCZ support this possibility. Intriguingly, antipsychotic medication has been reported to reduce neutrophil-to-lymphocyte ratio in SCZ ^91^. Elucidation of the interactions of circulating immune cells with astrocytes, aside from BBB integrity effects, in the SCZ context may reveal potential therapeutic targets.

In conclusion, we propose that an intrinsic neuroinflammatory imbalance affecting astrocytes that is evidenced by NF-κB dysfunction and elevated cytokine secretion may underlie, at least in part, the developmental disturbances that shape brain changes in SCZ pathogenesis. We also suggest that astrocyte-derived IL-8 has the potential to affect the cerebrovascular net, thus disrupting its microenvironment. The immune dysregulation of astrocytes should be considered as a target in the development of new drug therapies for SCZ.

## Author contributions

PT, JMN, and SKR conceived and designed the study. PT performed all iPSC-derived cell experiments. JMN performed proteomic experiments, analyses, and interpretation. BSC and TM performed CAM experiments, analyses, and interpreted data. SD performed and analyzed RT-qPCR experiments. JG, CTR, JCFM, and DPG performed and analyzed cytokine secretome data. LOP analyzed the data and discussed the results. VP contributed to angiogenesis experiments, analyses, and data interpretation. DMS supervised proteomic experiments and data interpretation. PT and JMN interpreted data, wrote, prepared figures, and revised the manuscript. SKR coordinated the study. All authors revised and contributed to the final version of this manuscript.

## Acknowledgments

We acknowledge the contributions on technical support of Fernanda Albuquerque, Gabriela Lopes Vitória, Ismael Carlos da Silva Gomes, Jarek Sochacki, Paulo Baldasso, Mariana Fioramonte, Marcelo do Nascimento Costa and Renata Maciel Santos. This project was sponsored by Fundação de Amparo à Pesquisa do Estado do Rio de Janeiro (FAPERJ), Conselho Nacional de Desenvolvimento Científico e Tecnológico (CNPq), Coordenação de Aperfeiçoamento de Pessoal de Nível Superior (CAPES), Instituto Nacional de Neurociência Translacional (INNT), Banco Nacional de Desenvolvimento (BNDES), the São Paulo Research Foundation (FAPESP) #2014/21035-0 (JMN), 2017/25588-1 (DMS) and 2019/00098-7 (DMS) and ANID/FONDECYT # 1190083 (VP), CONICYT/ANID Fellowships for PhD #21150781 (BC), in addition to intramural grants from D’Or Institute for Research and Education.

---

**Supplementary Table 1. Summary of hiPSC characteristics**

**Supplementary Table 2. Proteomics data of whole-cell hiPSC-derived astrocytes from control and schizophrenia.**

**Supplementary Table 3. Proteomics data of conditioned media of hiPSC-derived astrocytes from control and schizophrenia stimulated or not with TNF-α.**

