## Supplementary Table 1. for "Induced pluripotent stem cell-derived astrocytes from patients with schizophrenia exhibit an inflammatory phenotype that affects vascularization"

| Cell ID | Cell Line | Gender | Age of | Cell Source | Institution | Previous Publications | Use in experiments |
|---|---|---|---|---|---|---|---|
| GM23279A | CTRL 1 | F | 20 | Dermal fibroblasts | Coriell Biobank | Casas et al. (2018), Ledur et al. (2020), | Figures 2-7 and Supplementary |
| CF1 | CTRL 2 | M | 37 | Dermal fibroblasts | D'Or Institute for Research | Casas et al. (2018), Ledur et al. (2020), | Figures 2-7 and Supplementary |
| CF2 | CTRL 3 | M | 31 | Dermal fibroblasts | D'Or Institute for Research | Casas et al. (2018), Ledur et al. (2020), | Figures 2-7 and Supplementary |
| C15 | CTRL 4 | F | 16 | Urine cells | D'Or Institute for Research | Ledur et al. (2020), Trindade et al. (2020) | Figures 3-7 and Supplementary |
| GM23760B | SCZ 1 | M | 26 | Dermal fibroblasts | Coriell Biobank | Casas et al. (2018) | Figures 2-7 and Supplementary |
| GM23761B | SCZ 2 | F | 27 | Dermal fibroblasts | Coriell Biobank | Casas et al. (2018) | Figures 2-7 and Supplementary |
| EZQ4 | SCZ 3 | M | 42 | Dermal fibroblasts | D'Or Institute for Research | Casas et al. (2018) | Figures 2-7 and Supplementary |
